# The orchestrated cellular and molecular responses of the kidney to endotoxin define the sepsis timeline

**DOI:** 10.1101/2020.05.27.118620

**Authors:** Danielle Janosevic, Jered Myslinski, Thomas McCarthy, Amy Zollman, Farooq Syed, Xiaoling Xuei, Hongyu Gao, Yunlong Liu, Kimberly S. Collins, Ying-Hua Cheng, Seth Winfree, Tarek M. El-Achkar, Bernhard Maier, Ricardo Melo Ferreira, Michael T. Eadon, Takashi Hato, Pierre C. Dagher

## Abstract

Clinical sepsis is a highly dynamic state that progresses at variable rates and has life-threatening consequences. Staging patients along the sepsis timeline requires a thorough knowledge of the evolution of cellular and molecular events at the tissue level. Here, we investigated the kidney, an organ central to the pathophysiology of sepsis. Single cell RNA sequencing revealed the involvement of various cell populations in injury and repair to be temporally organized and highly orchestrated. We identified key changes in gene expression that altered cellular functions and can explain features of clinical sepsis. These changes converged towards a remarkable global cell-cell communication failure and organ shutdown at a well-defined point in the sepsis timeline. Importantly, this time point was also a transition towards the emergence of recovery pathways. This rigorous spatial and temporal definition of murine sepsis will uncover precise biomarkers and targets that can help stage and treat human sepsis.

## Introduction

Acute kidney injury (AKI) is a common complication of sepsis that doubles the mortality risk. In addition to failed homeostasis, kidney injury can contribute to multi-organ dysfunction through distant effects. Indeed, the injured kidney is a significant mediator of inflammatory chemokines, cytokines, and reactive oxygen species that can have both local as well as remote deleterious effects ^1-4^. Therefore, understanding the complex pathophysiology of kidney injury is crucial for the comprehensive treatment of sepsis and its complications.

We have recently shown that renal injury in sepsis progresses through multiple phases. These include an early inflammatory burst followed by a broad antiviral response and culminating in translation shutdown and organ failure ^5^. In a non-lethal and reversible model of endotoxemia, organ failure was followed by spontaneous recovery. The exact cellular and molecular contributors to this multifaceted response remain unknown. Indeed, the kidney is architecturally a highly complex organ in which epithelial, endothelial, immune and stromal cells are at constant interplay. Therefore, we now examined the spatial and temporal progression of endotoxin injury to the kidney using single cell RNA sequencing (scRNAseq). Our data revealed that cell-cell communication failure is a major contributor to organ dysfunction in sepsis. Remarkably, this phase of communication failure was also a transition point where recovery pathways were activated. We believe this spatially and temporally anchored approach to sepsis pathophysiology is crucial for identifying potential biomarkers and therapeutic targets.

## Results

### Single cell RNA sequencing and spatial transcriptomics identify and localize known and novel renal cell populations

We harvested a cumulative amount of 63,287 renal cells obtained at 0, 1, 4, 16, 27, 36 and 48 hours after endotoxin (LPS) administration. The majority of renal epithelial, immune and endothelial cell types were represented (**Fig. 1a**). Note the absence of podocyte and mesangial cells, which can be a limitation of single cell RNAseq renal dissociation procedures ^6^. Cluster identities were assigned and grouped using known classical phenotypic markers **(Fig. 1b, Supplementary Fig. 1a)** ^7-11^. Interestingly, the UMAP-based computational layout of epithelial clusters recapitulated the normal tubular segmental order in the nephron. This indicates that gene expression gradually changes among neighboring tubular segments along the nephron. Note that the expression of cluster-defining markers varied significantly during the injury and recovery phases of sepsis (**Fig. S1b; Supplementary Table 1**). Therefore, we also identified a set of genes that are conserved across time for a given cell type (**Fig. S1c**).

**Fig. 1:**
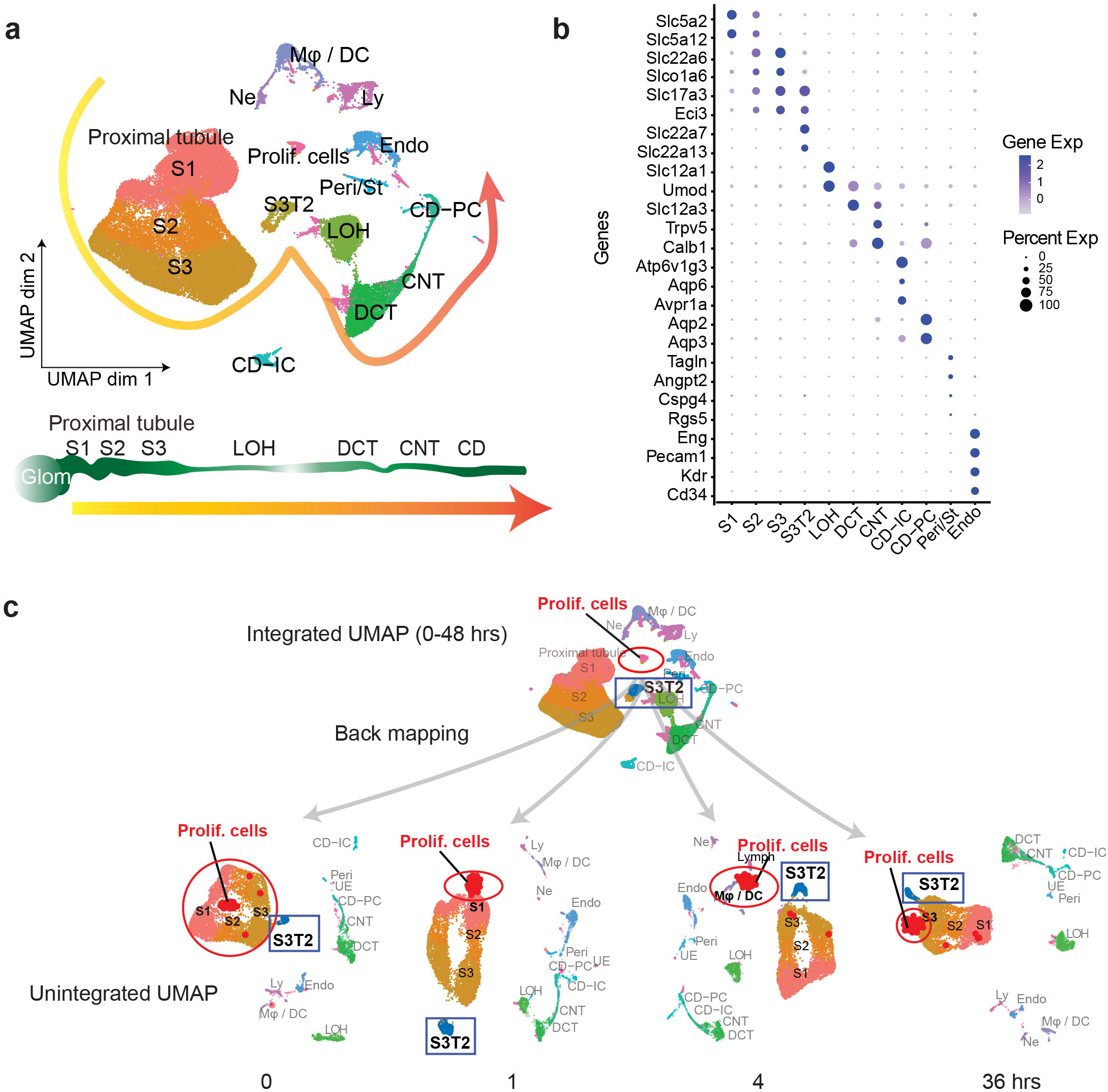
Single cell RNA sequencing identifies renal epithelial, endothelial, stromal and immune cell types in the murine septic kidney. **a** Integrated UMAP of kidney cell clusters from control and LPS-treated mice (0, 1, 4, 16, 27, 36 and 48 hours after LPS injection). Actual anatomical layout of kidney nephronal segments is shown below UMAP. **b** Dot plot of representative genes defining indicated cell types. **c** Back mapping of cells from the integrated UMAP onto unintegrated UMAPs of select time points. Highlighted are the proliferating cell cluster (red circle) and S3T2 cluster (blue box). CD, collecting duct. CD-IC, collecting duct-intercalated cells. CD-PC, collecting duct-principle cells. CNT, connecting tubule. OCT, distal convoluted tubule. Endo, endothelial cells. Exp, expression. Glom, glomerulus. LOH, Loop of Henle. LPS, endotoxin. Ly, lymphocytes. Mφ-DC, macrophage-dendritic cells. Ne, neutrophil. Peri/St, mixed pericyte and stromal cells. Prolif. Cells, proliferative cells. PT, proximal tubule. S1, first segment of PT. S2, second segment of PT. S3, third segmaent of PT. S3T2, S3 type 2 cells.

In the integrated UMAP (**Fig. 1a**), we noted the presence of a proliferative cell cluster (*Cdk1* and *Ki67* expression). By back mapping to time-specific unintegrated UMAPs, we determined that these proliferating cells could be traced to specific cell types at various points along the sepsis timeline (**Fig. 1c**). For example, within the first hour after LPS, these proliferative indices were expressed primarily in S1 cells. These cells are the site of LPS uptake in the kidney as we have previously shown ^12-14^. At later time points, proliferative indices are seen in macrophages (4 hours) and S3 cells (36 hours) (**Fig. 1c**). These proliferative indices reflect cell cycle activity which may be involved in injury, repair or recovery processes ^15^.

We also noted the presence of a proximal tubular cluster expressing unique gene identifiers: *Agt, Rnf24, Slc22a7* and *Slc22a13* (**Fig. 2a**). This is likely the proximal tubular S3-Type 2 (S3T2) reported by others ^16^. This cluster maintained a separate and distinct identity throughout the sepsis timeline (**Fig. 1c**). Because the location of S3T2 is currently unknown, we performed in-situ spatial transcriptomics on septic mouse kidneys ^17^. We then integrated our scRNAseq with the in-situ RNAseq in order to map our scRNAseq clusters onto the tissue (**Supplementary Fig. 2a, S2b**). We found that the classical S3 cluster localizes to the cortex while S3T2 is in the outer stripe of the outer medulla (**Fig. 2b, Supplementary Fig. 2b**). We confirmed the location of S3T2 to the OS-OM with single molecular FISH (**Supplementary Fig. 2c**). The differential gene expression between S3 and S3T2 is likely dictated by regional differences in the microenvironments of the cortex and the outer stripe.

**Fig. 2:**
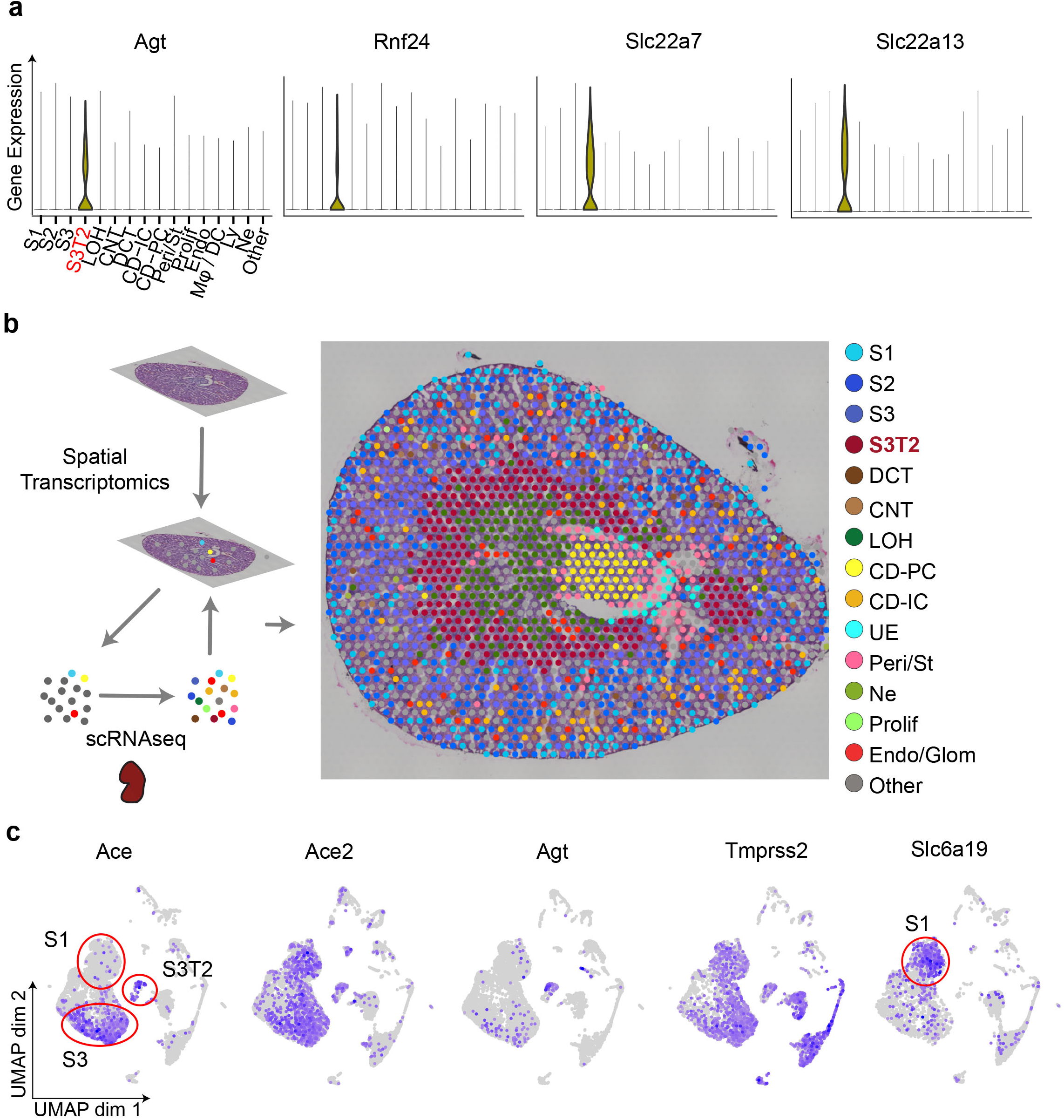
Spatial transcriptomics localize S3-Type 2 cells to the outer stripe of the outer medulla. **a** Violin plots of S3T2 defining markers. **b** Integration of spatial transcriptomics and scRNAseq. Spatial transcriptomics were performed on a slice of mouse kidney after cecal ligation and puncture. This yielded 7 clusters that were expanded to 15 cell types by integrating spatial transcriptomics with scRNAseq data from LPS-treated mice. See also **Supplemental Fig. 2b. c** Feature plots of select renin-angiotensin system and other SARS-CoV-2-related genes. See also **Supplemental Fig. 3**. CD, collecting duct. CD-IC, collecting duct-intercalated cells. CD-PC, collecting duct-principle cells. CNT, connecting tubule. OCT, distal convoluted tubule. Endo, endothelial cells. Exp, expression. Glom, glomerulus. Hrs, hours. LOH, Loop of Henle. Ly, lymphocytes. Mφ-DC, macrophage-dendritic cells. Ne, neutrophil. Peri/St, mixed pericyte and stromal cells. Prolif., proliferative cells. PT, proximal tubule. S1, first segment of PT. S2, second segment of PT. S3, third segment of PT. S3T2, S3 type 2 cells. scRNAseq, single cell RNA sequencing.

Because angiotensinogen (*Agt*) was strongly expressed in S3T2, we examined the expression of other components of the renin-angiotensin system (RAS). We first noted the absence of *Ace* expression in S1 tubular cells (**Fig. 2c, Supplementary Fig. 3**). In contrast, *Ace2* was strongly expressed in S1, S3 and S3T2 cells. There is currently great interest in understanding the biology of *Ace2* because of its role in SARS-CoV-2 cellular invasion. Other essential components of the SARS-CoV-2 entry mechanism include *Tmprss2* and *Slc6a19* ^18-22^. While *Tmprss2* was expressed in all proximal tubular segments, *Slc6a19* was more strongly expressed in S1 throughout the sepsis timeline. This may point to the S1 tubular segment as one point of entry of SARS-CoV-2 into the kidney.

### Cell trajectory and velocity field analyses of scRNAseq characterize subpopulations of immune cells

The immune cell profile in the septic kidney was time-dependent and showed a five-fold increase in immune cells, primarily macrophages (**Fig. 3a, 3b**). We noted two distinct macrophage clusters denoted as Macrophage A and Macrophage B (Mφ-A, Mφ-B). Both of these clusters expressed classical macrophage markers such as Cd11b (*Itgam*) (**Fig. 3c**). However, they differed in the expression of *Adgre1* (F4/80, Mφ-A) and *Ccr2* (Mφ-B). The accumulated macrophages were predominantly Mφ-A. We noted the absence of proliferation markers (*Cdk1, Ki67*) in this cluster, raising the possibility that this may be an infiltrative macrophage type (**Fig. 3d**). The Mφ-B cluster, located between Mφ-A and conventional dendritic cells (cDC) expressed also cDC markers such as MHC-II subunit genes (*H2-Ab1*) and Cd11c (*Itgax*) indicating that it is an intermediary macrophage type (**Fig. 3c**). This continuum between macrophages and dendritic cells in the kidney has been reported ^23-26^. Interestingly, Mφ-B cells expressed proliferation markers (*Cdk1, Ki67*) and thus, may be differentiating towards a Mφ-A or cDC phenotype (**Fig. 3c**). Pseudotime and velocity field analysis suggested that at earlier time points (1 hour) Mφ-B was differentiating toward Mφ-A phenotype. At later time points (36 hours) the velocity field suggested that Mφ-B was differentiating towards cDC but pseudotime analysis was inconclusive (**Fig. 3e**). Similarly, the Mφ-A cluster also showed two subclusters on the RNA velocity map (**Supplementary Fig. 4a**). One of the subclusters showed increased expression of alternatively activated macrophages (M2) markers such as *Arg1* (Arginase 1) and *Mrc1* (Cd206) ^27^ at later time points (36 hours, **Supplementary Fig. 4b**). Therefore, RNA velocity analysis may be a useful tool in distinguishing macrophage subtypes in scRNAseq data.

**Fig. 3:**
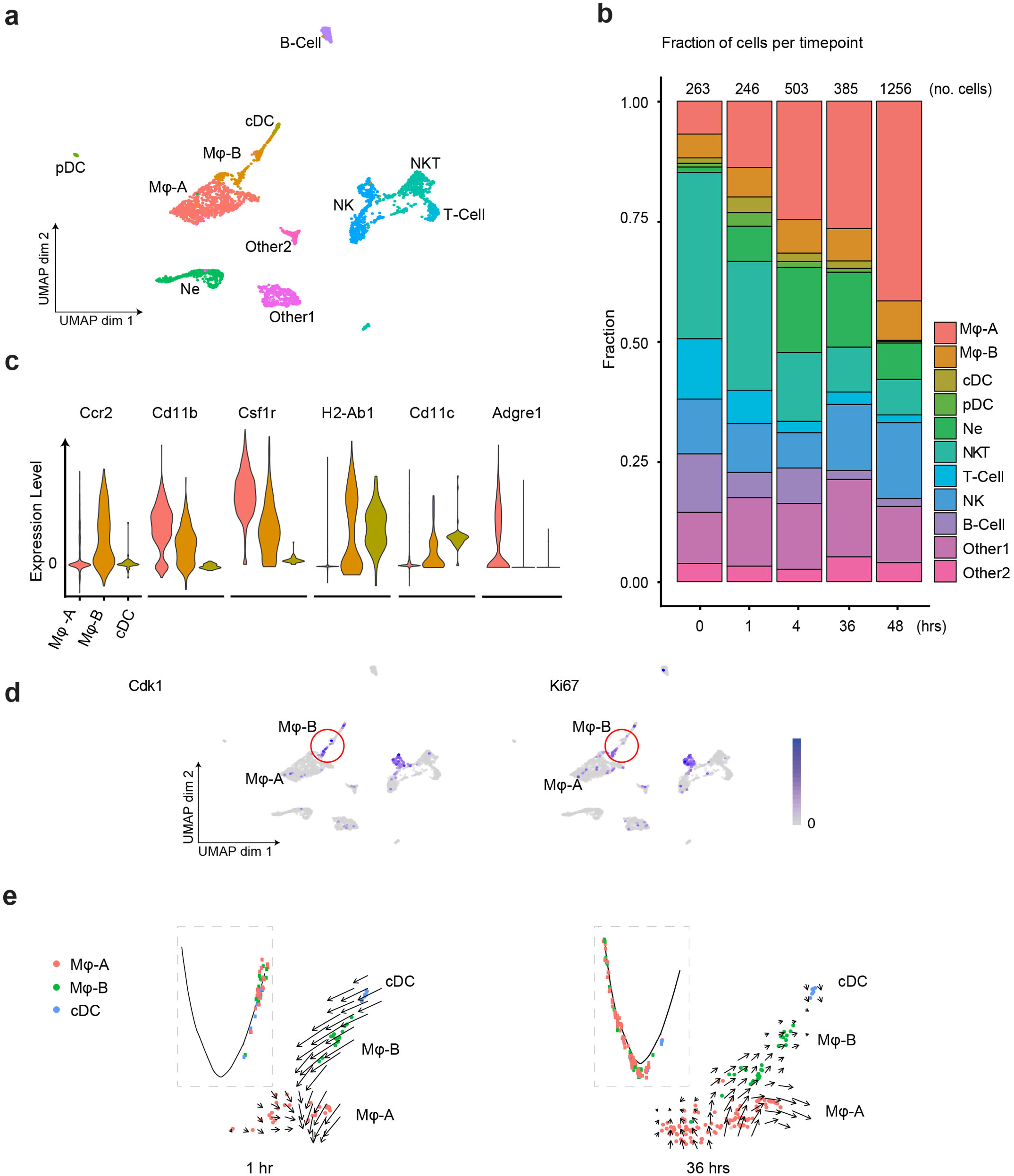
Sepsis induces dynamic changes in renal immune cell composition, pseudotime states and RNA velocity. **a** Integrated UMAP of the immune cell clusters from control and LPS-treated mice (0, 1, 4, 16, 27, 36 and 48 hours after LPS injection). Other 1 and Other 2 are Cd45+ cells with mixed epithelial and immune markers. b Stacked bar plot with fractions of immune cells (relative to total number of cells) shown in the y-axis, at 0, 1, 4, 36 and 48 hours after LPS. The total number of immune cells is indicated at the top of the bar for each time point. **c** Integrated violin plots from all time points for indicated genes defining subtypes of macrophages and DCs are shown. d Feature plots of proliferation markers expression from integrated time points in the immune cell subsets. **e** Integrated cell trajectory analyses and RNA velocity fields for macrophages and dendritic cells shown at indicated time points. cDC, conventional dendritic cell. Hrs, hours. Mφ-A, macrophage-A. Mφ-B, macrophage-B. Ne, neutrophil. NK, natural killer cells. NKT, natural killer T-cells. pDC, plasmacytoid dendritic cell. T-cell, Cd3+ T-lymphocytes.

In T-cells, while *Cd4* expression was minimal at all time points, the expression of *Cd8* was robust and relatively preserved over time (**Fig. S4c)**. We also noted an increase of a distinct plasmacytoid dendritic cell cluster at one hour (pDC). These pDCs, along with natural killer (NK) cells, are known to signal through the interferon-gamma pathway and stimulate Cd8 expression ^28,29^. This supports the early antiviral response we have previously reported in this sepsis model ^5^.

### Cell trajectory and velocity field analyses of scRNAseq characterize subpopulations of epithelial and endothelial cells

We next examined the phenotypic changes in various cell populations along the sepsis timeline. At each time point in sepsis, cells exhibited various states of gene expression that are well defined with pseudotime analysis. We note that at any given time point, directional progression of states along pseudotime correlated well with real time state changes (**Fig. 4a**). Note that the endothelium exhibited changes in states as early as 1 hour, while S1 showed changes at later time points (4 hours). These sequential state changes may reflect the spatial and temporal propagation of LPS signaling in the kidney. As sepsis progressed, many cell types lost function-defining markers while acquiring novel ones. For example, S1 and S3 lost classical markers like *Slc5a2* (SGLT2) and *Aqp1* and expressed new genes involved in antigen presentation such as *H2-Ab1* (MHC-II) and *Cd74* (**Fig. 4b**). Moreover, the highly distinct phenotypes that differentiated S1 from S2/S3 at baseline merged into one phenotype for all three sub-segments by 16 hours after LPS (**Fig. 4c**). However, despite the apparent convergent phenotype at 16 hours, additional analytical approaches such as RNA velocity revealed significant differences in RNA splicing kinetics between S1 and S3 segments at this time point. In addition, RNA velocity revealed the presence of two subclusters within the S3 segment at 16 hours (**Fig. 4d**). These two velocity subclusters did not correlate with the two states seen in pseudotime analysis. This indicates that multiple analytic approaches are needed to fully characterize cellular changes along the sepsis timeline.

**Fig. 4:**
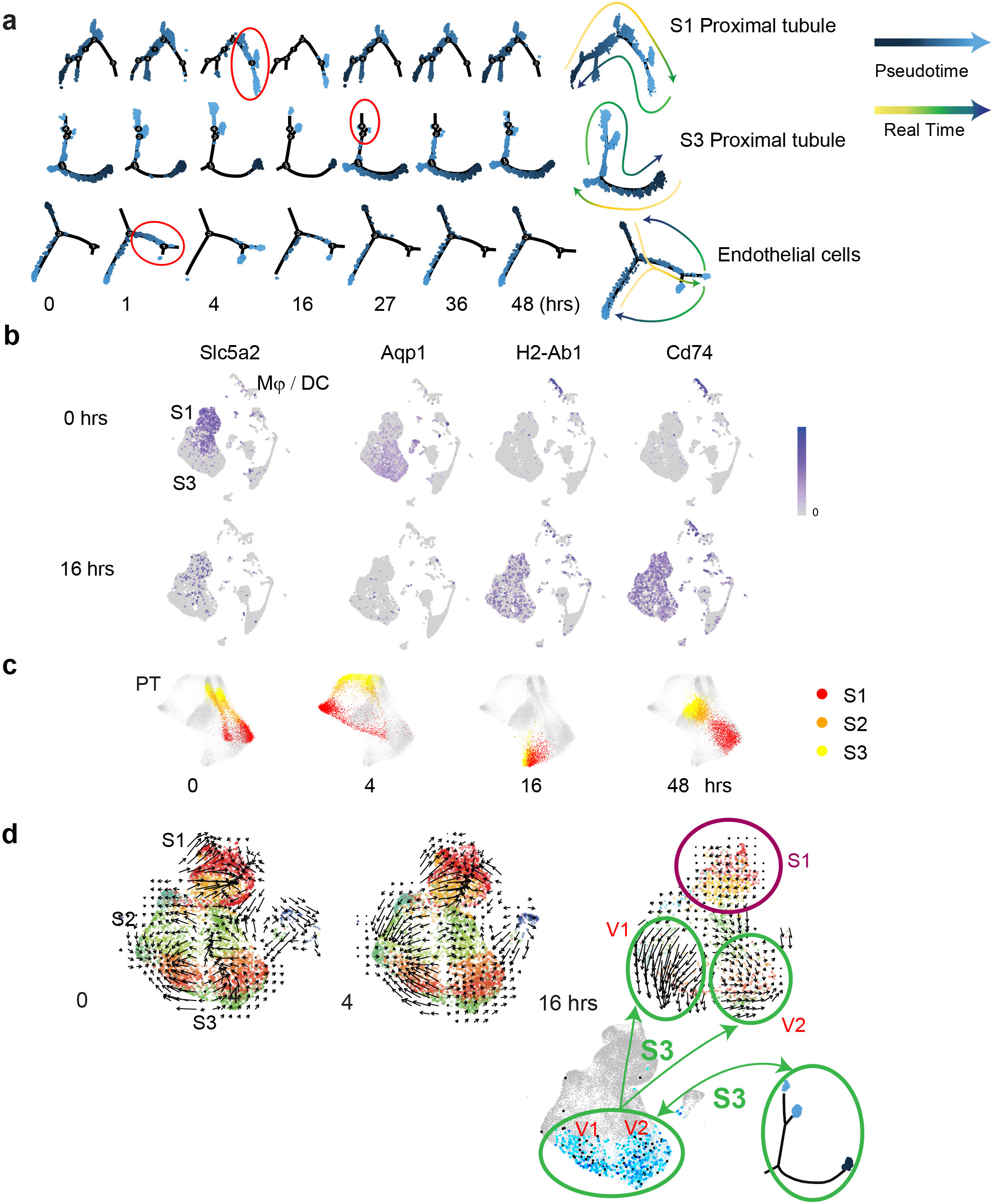
Sepsis alters pseudotime states, phenotypic gene expression and RNA velocity in renal cell populations. **a** Cell trajectory analysis for S1, S3 and endothelial cells shown at indicated time points. Highlighted in red circles are significant state transitions in respective cell types. The last cell trajectory shown for each cell type is integrated from all time points. It highlights the correspondence between pseudotime and real time. b Feature plots of select genes shown at indicated time points highlighting proximal tubular phenotypic changes. **c** Time-specific S1, S2 and S3 PT cells (red, orange, yellow) overlaid on composite t-SNE map of all PT cells (grey). d RNA velocity fields for S1, S2, and S3 proximal tubular cells are shown at indicated time points. Two velocity subfields V1 and V2 in S3 cells are circled in green. Projections of two pseudotime S3 states (light blue, dark blue dots) onto the S3 velocity fields do not show a 1:1 correspondence with the two velocity subfields V1 and V2. Hrs, hours. Mφ-DC, macrophage-dendritic cells. Ne, neutrophil. PT, proximal tubule. S1, first segment of PT. S2, second segment of PT. S3, third segment of PT. S3T2, S3 type 2 cells. V1, velocity subfield 1. V2, velocity subfield 2.

### Sepsis induces time and cell-specific genes and pathways

We next show gene expression profiles in select cell types along the sepsis timeline. In this analysis, we included endothelial cells, pericyte/stromal cells, macrophages and S1 tubular cells. Within 1 hour of LPS exposure, most cell types showed decreased expression of select genes involved in ribosomal function, translation and mitochondrial processes such as *Eef2* and *Rpl* genes (**Fig. 5a, Supplementary Fig. 5a**). This reduction peaked at 16 hours and recovered by 27 hours. Concomitantly, most cell types exhibited increased expression of several genes involved in inflammatory and antiviral responses such as *Tnfsf9, Cxcl1, Ifit1*, and *Irf7*. However, this increase was not synchronized among all cell populations. Indeed, it occurred as early as 1 hour in endothelial cells, macrophages and pericyte/stromal cells, all acting as first responders. In contrast, epithelial cells were late responders, with increases in inflammatory and antiviral responses occurring between 4 and 16 hours. In fact, four hours after LPS administration, cluster-specific GO terms were indistinguishable among the majority of cell types with enrichment in terms related to defense, immune and bacterium responses (**Fig. 5b**). One noted exception was the S3T2 cells (outer stripe S3) which did not enrich as robustly as other cell types in these terms. It mostly maintained an expression profile related to ribosomes, translation and drug transport throughout the sepsis timeline (**Supplementary Fig. 6**). Other players of interest in sepsis pathophysiology such as prostaglandin and coagulation factors are described in **Supplementary Figure 5b**.

**Fig. 5:**
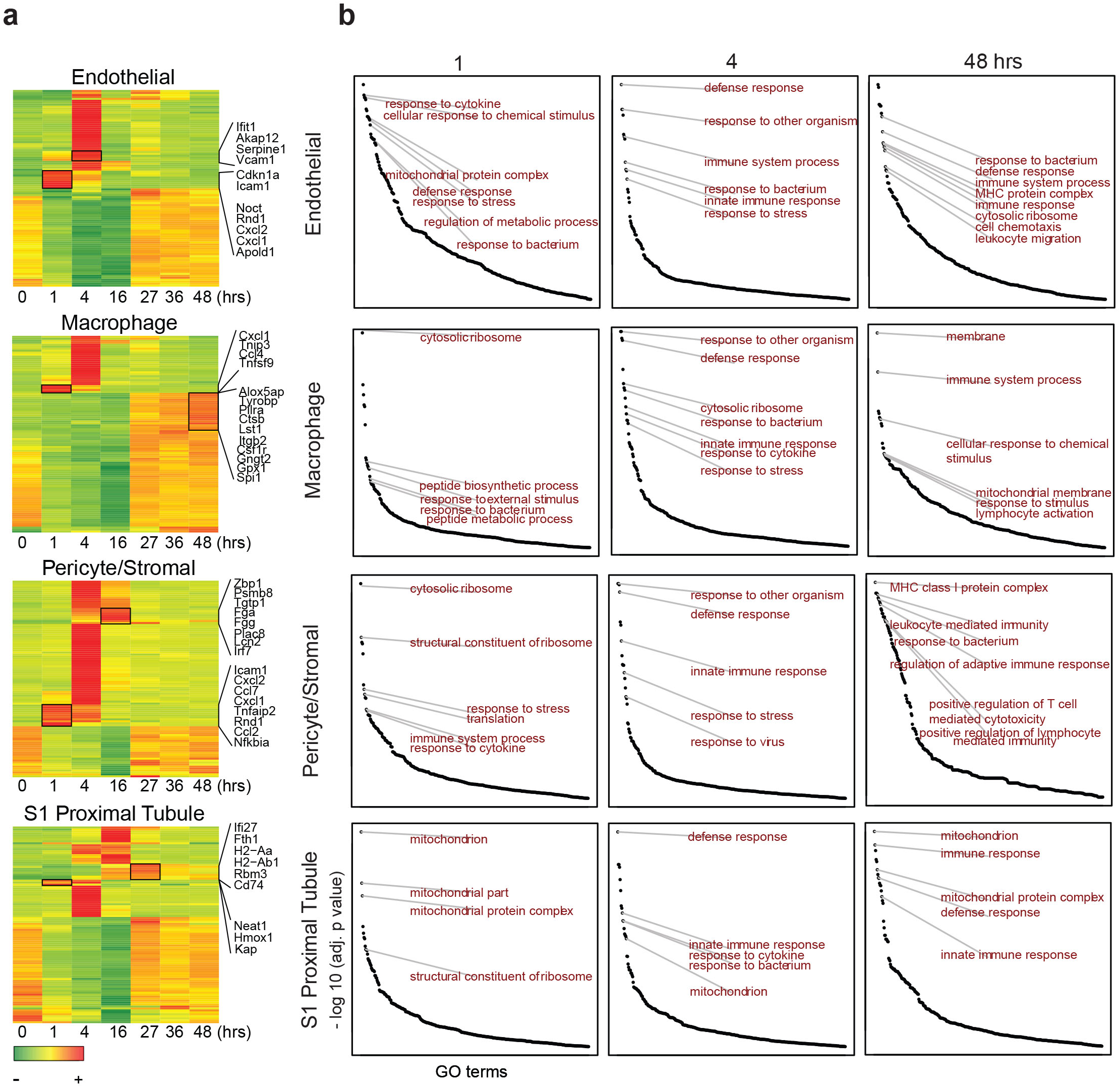
Sepsis induces time and cell-specific genes and pathways. **a** Heatmaps of select cell types with top 100 differentially expressed genes across the sepsis timeline (0-48 hours). Select genes are shown for each cell type. **b** Time dependent enrichment of gene ontology terms for indicated cell types. GO terms are sorted in order of statistical significance. Hrs, hours. GO, gene ontology biological processes

At the 48-hour time point, while S1 cells partially recovered to baseline, the macrophages showed increased expression of genes involved in phagocytosis, cell motility and leukotrienes, broadly representative of activated macrophages (e.g. *Csf1r, Lst1, Capzb, S100a4, Cotl1, Alox5ap*, **Fig. 5a**). Intriguingly, at this late time point, the pericyte/stromal cells are enriched in unique terms related to specific leukocyte and immune cell types such as lymphocyte-mediated immunity, T cell mediated cytotoxicity and antigen processing and presentation. This suggests that the pericyte may function as a transducer between epithelia and other immune cells.

### Sepsis alters cell-cell communication in the murine kidney

Therefore, we next examined comprehensively cell-cell communication along the sepsis timeline. We show select examples of cell type-specific receptor ligand pairs. For example, we found that S1 and endothelial cells communicate with the *Angpt1* (Angiopoetin 1) and *Tek* (Tie2) ligand-receptor pair at baseline and throughout the sepsis timeline (**Fig. 6a-b, Supplementary Fig. 7a**). In contrast, *C3* was strongly expressed in pericyte/stromal cells, while its receptor *C3ar1* localized to macrophage/DCs. This communication, present at baseline, did increase along the sepsis timeline with additional players such as S1 participating in the cross talk (**Supplementary Fig. 7**). Another strong communication was noted between endothelial cells and macrophage/lymphocytes using the *Ccl2* and *Ccr2* receptor-ligand pair. The architectural layout of these four cell types, with pericytes and endothelial cells residing between proximal tubule and macrophage/DCs may explain these complex communication patterns ^6^. Such communication patterns among these four cell types may also explain macrophage clustering around S1 tubules at later time points in sepsis as we previously reported ^13^.

**Fig. 6:**
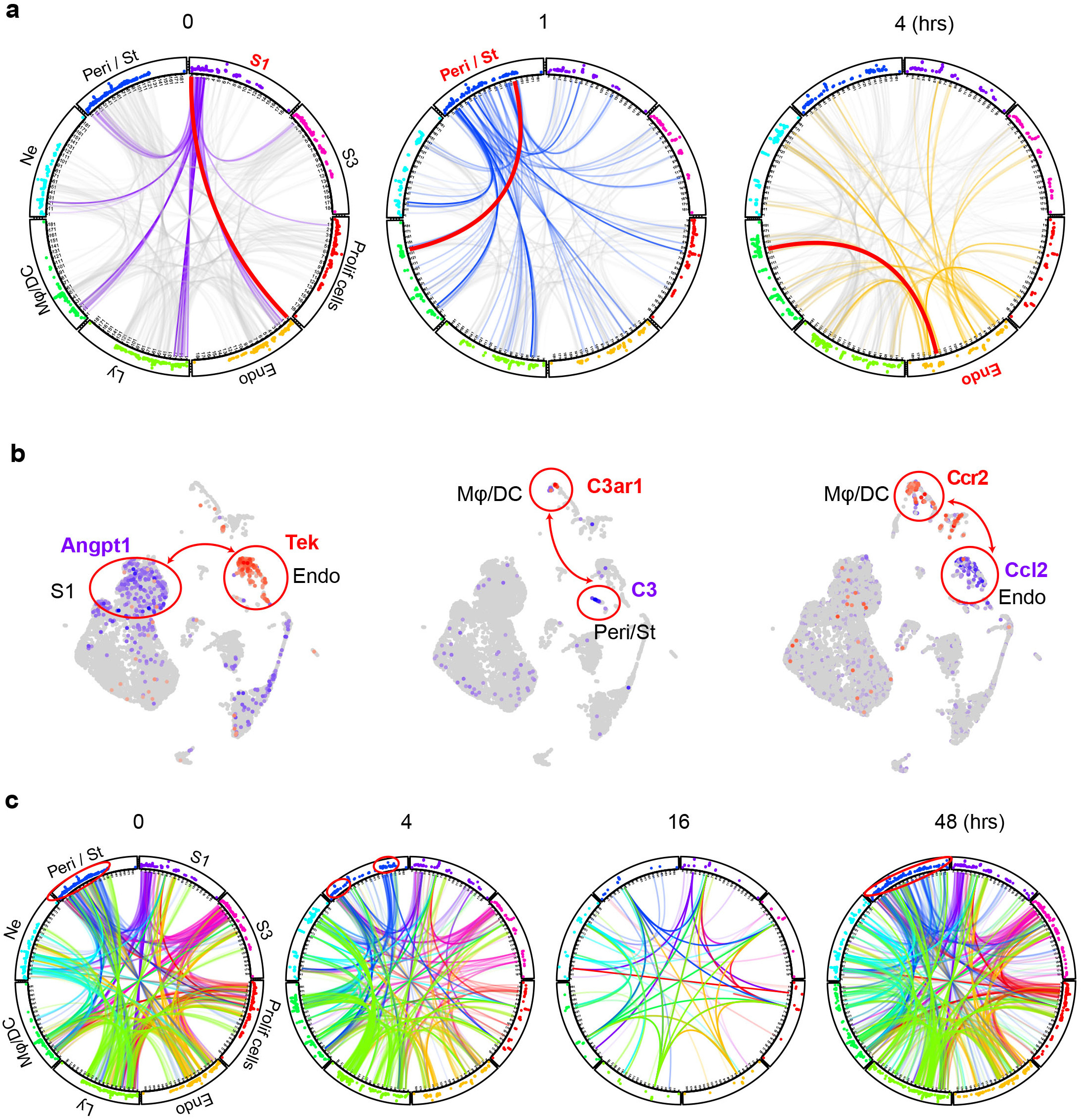
Sepsis alters cell-cell communication in the murine kidney. **a** Receptor-ligand pairs for indicated cell types are displayed in circular plots. The data was generated using the CellPhone database. For clarity, communication between one cell type and all others is shown (purple lines: 0 hr for S1, blue lines: 1 hr for Peri/St and yellow lines: 4 hrs for endothelial cells). Other cell-cell communications in each circular plot are shown in light grey in the background. In each circular plot, the red line connects the specific receptor-ligand pair highlighted in panel B. Dots in the outer track of the circle represent specific ligands or receptors and are positioned identically for all cell types. The height of dots correlates with statistical significance (all dots are less than adjusted p.value <0.05). The identity of each dot is given in **Supplementary Table 2**. **b** Feature plots of receptor-ligand pairs between specified cell types as highlighted by the red line in panel A. In each feature plot, the ligand is shown in purple and the receptor in red. **c** Circular plots displaying receptor-ligand interactions between all cell types at specified time points. Examples of change in communication patterns are shown in the red circles in the outer track of the plot at 0, 4 and 48 hours. Note the dramatic drop in cell communication at 16 hours. Endo, endothelial cells. Hrs, hours. Ly, lymphocytes. Mφ-DC, macrophage-dendritic cells. Ne, neutrophil. Peri/St, mixed pericyte and stromal cells. Prolif. Cells, proliferative cells. PT, proximal tubule. S1, first segment of PT. S3, third segment of PT.

When examined comprehensively, receptor-ligand signaling progressed from a broad pattern at baseline into a more discrete and specialized one 4 hours after LPS (**Fig. 6c, Supplementary Fig. 7b-c**). Sixteen hours after LPS, we noted a dramatic drop in cell-cell communication between all cell types. This communication failure may contribute to the transcription and translation shutdown we recently reported at this time point ^5^. In our reversible sepsis model, cell-cell communication recovered by 27-48 hours.

### Sepsis induces time and cell-specific changes in regulons

Transcription factors and their downstream targets (regulons) are important regulators of a myriad of pathways involved in the pathophysiology of sepsis. Therefore, we next examined the activity of regulons along the sepsis timeline in all renal cells. Surprisingly, we noted in many cell types an increase in regulon activity of key transcription factors at the 16-hour time point (**Supplementary Table 3**). As discussed above, this time point corresponds to translation shutdown as well as cell-cell communication failure. In S1, many of the regulons active at this time point are involved in cell differentiation, development, transcription and proliferation (*Sox4, Hoxb7, Srf*, **Fig. 7a-c**). Therefore, this 16-hour time point is not merely a time of complete shutdown and failure of the kidney. Rather, it is also a crucial transition point where key regulators of recovery and healing are being activated.

**Fig. 7:**
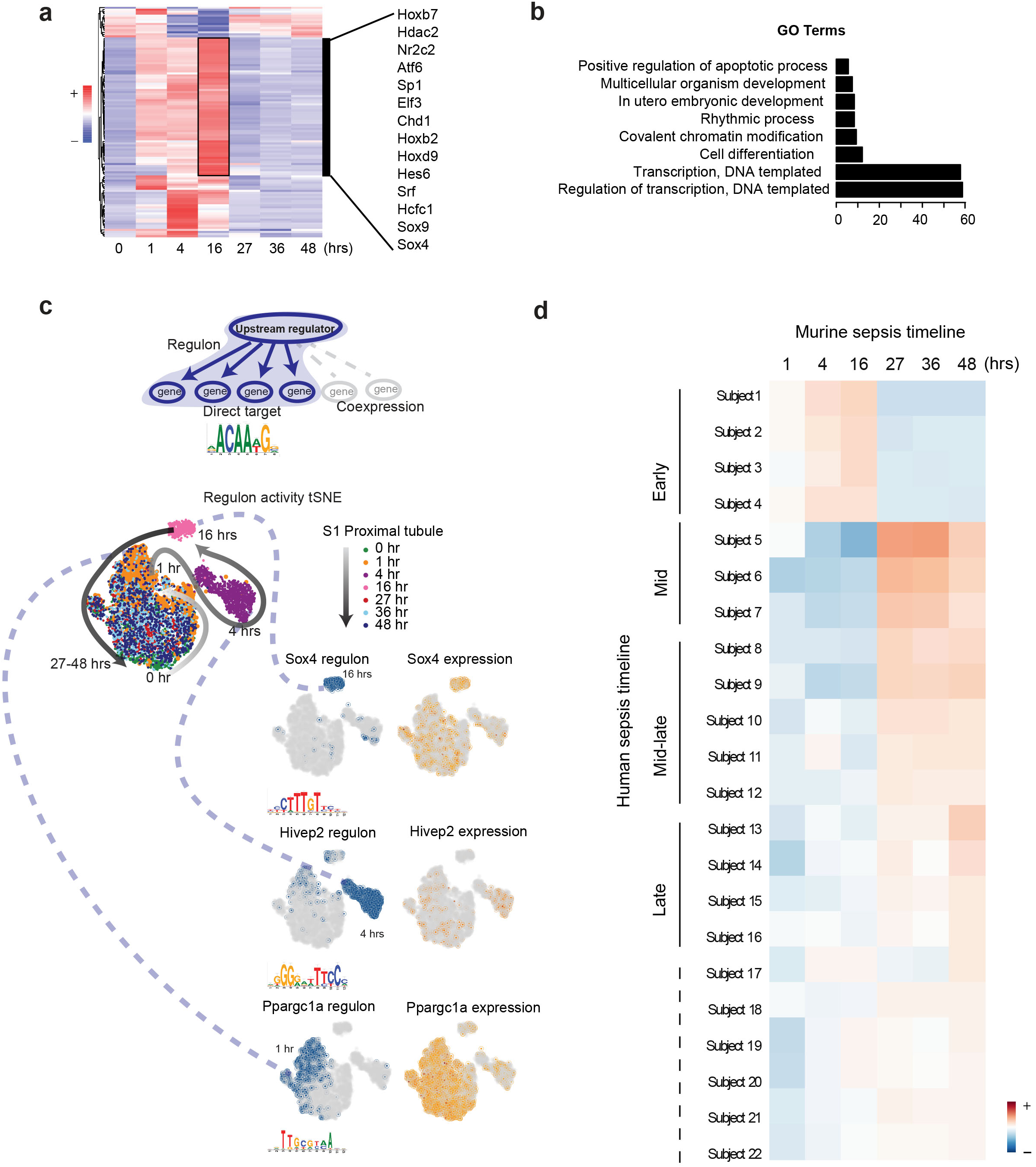
Sepsis induces time and cell-specific changes in regulons. **a** SCENIC-derived heatmap of regulons for S1 tubules. Highlighted are select transcription factors with active regulons at the 16 hour time point. **b** Gene ontology pathway enrichment analysis derived from all regulons active at the 16-hour time point (Tables S4, SS). **c** t-SNE of proximal tubule S1 time-specific regulon activity. Select transcription factor expression (orange) and its corresponding regulon expression (blue) are shown. As shown for Sox4, note the temporal differences between the expression of the transcription factor itself and its regulon. **d** Heatmap of human sepsis kidney samples stratified based on aggregates of murine time-specific orthologues. The color scale indicates the degree of correlation based on Spearman’s ρ. Hr(s), hour(s). GO, gene ontology biological processes.

### The murine sepsis timeline allows staging of human sepsis

Finally, we asked whether our mouse sepsis timeline could be used to stratify human sepsis AKI. To this end, we selected the differentially expressed genes from all cells combined (pseudo bulk) for each time point across the mouse sepsis timeline (**Supplementary Table 4**). We then examined the orthologues of these defining genes in human kidney biopsies of patients with sepsis and AKI. The clinical data associated with these human biopsies did not allow further stratification or staging of the sepsis timeline (**Supplementary Table 5**). As shown in **Figure 7d**, our approach using the mouse data succeeded in partially stratifying the human biopsies into early, mid and late sepsis-related AKI. These findings suggest that underlying injury mechanisms are conserved, and the mouse timeline may be valuable in staging and defining biomarkers and therapeutics in human sepsis.

## Discussion

In this work, we provide comprehensive transcriptomic profiling of the kidney in a murine sepsis model. To our knowledge, this is the first description of spatial and temporal transcriptomic changes in the septic kidney that extend from early injury well into the recovery phase. Our data cover nearly all renal cell types and are time-anchored, thus providing a detailed and precise view of the evolution of sepsis in the kidney at the cellular and molecular level.

Using a combination of analytical approaches, we identified marked phenotypic changes in multiple cell populations along the sepsis timeline. The proximal tubular S1 segment exhibited significant alterations consisting of early loss of traditional function-defining markers (e.g., SGLT2). Similar losses of function-defining markers along the nephron may explain the profound derangement in solute and fluid homeostasis seen in sepsis. Concomitantly, we observed novel epithelial expression of immune-related genes such as those involved in antigen presentation. This indicates a dramatic switch in epithelial function from transport and homeostasis to immunity and defense. These phenotypic changes were reversible, thus underscoring the remarkable resilience and plasticity of the renal epithelium.

In addition, our combined analytical tools clearly identified unique subclusters within each epithelial cell population (e.g., cortical S3 and OS S3). These subclusters likely represent novel populations that may be in part influenced by the complex microenvironments in the kidney. It is likely that such microenvironments define unique features in epithelial subpopulations such as the expression of complete SARS-CoV-2 machinery in S1.

Similarly, we also identified unique features in immune-cell populations. For example, the combined use of RNA velocity field and pseudotime analyses uncovered differences in macrophage subtypes relating to RNA kinetics and cell differentiation trajectories. Of note is that these subtypes only partially matched the traditional flow cytometry-based classification of macrophages (e.g., M1/M2). Therefore, the use of single-cell RNA seq is a powerful approach that will add to and complement our current understanding of the immune cell repertoire in the kidney.

Additional approaches such as receptor-ligand crosstalk and gene regulatory network analyses identified unique cell- and time-dependent players involved in sepsis pathophysiology. Importantly, the expression of genes involved in vectorial transport, inflammation, vascular health and cell-cell communication varied greatly along the sepsis timeline, and required simultaneous contributions from multiple cell types. However, these complex interactions collapsed at the 16-hour time point. This indeed is a remarkable time in the sepsis timeline that we have previously investigated in multiple models of murine sepsis. It is the time where profound translation failure and organ shutdown occur. Our current data point to massive cell-cell communication failure as a key feature of this time point. Surprisingly, it is also at this time point that reparative pathways started to emerge. It is thus an important and defining point in sepsis that may have significant clinical implications.

Our work points to the urgent need for defining a more accurate and precise timeline for human sepsis. Such definition will guide the development of biomarkers and therapies that are cell and time specific. We show evidence supporting the relevance of murine models and their usefulness in staging human sepsis. These precisely time- and space-anchored data will provide the community with rich and comprehensive foundations that will propel further investigations into human sepsis.

## Methods

### Experimental Model and Subject Details

#### Animal model

Male C57BL/6J mice were obtained from the Jackson Laboratory. Mice were 8-10 weeks of age and weighed 20-25 g. They were subjected to a single dose of 5 mg/kg LPS tail vein injection (E. coli serotype 0111:B4 Sigma). Animals were sacrificed at 0, 1, 4, 16, 27, 36 and 48 hours after LPS (both kidneys per animal for each time point). For spatial transcriptomics experiments, cecal ligation and puncture (CLP) was performed under isoflurane anesthesia; 75% of the mouse cecum was ligated and punctured twice with a 27-gauge needle and the mouse sacrificed and kidneys harvested 6 hours later.

#### Study approval

All animal protocols were approved by the Indiana University Institutional Animal Care Committee and conform to the NIH (*Guide for the Care and Use of Laboratory Animals*, National Academies Press, 2011). The study in humans was approved by the Indiana University Institutional Review Board (protocol no. 1601431846). As only archived human biopsies were used in this study, the Institutional Review Board determined that informed consent was not required.

### Isolation of single cell homogenate from murine kidneys

Murine kidneys were transported in RPMI1640 (Corning), on ice immediately after surgical procurement. Kidneys were rinsed with PBS (ThermoFisher) and minced into eight sections. Each sample was then enzymatically and mechanically digested with reagents from Multi-Tissue Dissociation Kit 2 and GentleMACS dissociator/tube rotator (Miltenyi Biotec). The samples were prepared per protocol “Dissociation of mouse kidney using the Multi Tissue Dissociation Kit 2” with the following modifications: After termination of the program “Multi_E_2”, we added 10 mL RPMI1640 (Corning) and 10% BSA (Sigma-Aldrich) to the mixture, filtered and homogenate was centrifuged (300 g for 5 minutes at 4°C). Cell pellet was resuspended in 1 mL of RBC lysis buffer (Sigma), incubated on ice for 3 minutes, and cell pellet washed three times (300 g for 5 minutes at 4°C). Annexin V dead cell removal was performed using magnetic bead separation after final wash, and the pellet resuspended in RPMI1640/BSA 0.04%. Viability and counts were assessed using Trypan blue (Gibco) and brought to a final concentration of 1 million cells/mL, exceeding 80% viability as specified by 10x Genomics processing platform.

### Single cell library preparation

The sample was targeted to 10,000 cell recovery and applied to a single cell master mix with lysis buffer and reverse transcription reagents, following the Chromium Single Cell 3’ Reagent Kits V3 User Guide, CG000183 Rev A (10X Genomics, Inc.). This was followed by cDNA synthesis and library preparation. All libraries were sequenced in Illumina NovaSeq6000 platform in paired-end mode (28bp + 91bp). Fifty thousand reads per cell were generated and 91% of the sequencing reads reached Q30 (99.9% base call accuracy). The total number of recovered cells for all time points was 63,287 cells, and per experiment was 9,191 (0 hour), 9,460 (1 hour), 9,865 (4 hours), 5,165 (16 hours), 7,678 (27 hours), 10,119 (36 hours), and 11,809 (48 hours after LPS).

### Single cell data processing

The 10x Genomics Cellranger (v. 2.1.0) pipeline was utilized to demultiplex raw base call files to FASTQ files and reads aligned to the mm10 murine genome using STAR ^30^. Cellranger computational output was then analyzed in R (v.3.5.0) using the Seurat package v. 3.0.0.9999, ^31^. Seurat objects were created for non-integrated and integrated (inclusive of all time points) using the following filtering metrics: gene counts were set between 200-3000 and mitochondrial gene percentages less than 50 to exclude doublets and poor quality cells. Gene counts were log transformed and scaled to 10^4^. The top 20 principle components were used to perform unsupervised clustering analysis, and visualized using UMAP dimensionality reduction (resolution 1.0). Using the Seurat package, annotation and grouping of clusters to cell type was performed manually by inspection of differentially expressed genes (DEGs) for each cluster, based on canonical marker genes in the literature ^8-10,32,33^. In some experiments, we used edgeR negative binomial regression to model gene counts and performed differential gene expression and pathway enrichment analyses (topKEGG, topGO, **Fig. 5, Supplementary Fig. 5a, Supplementary Fig. 6**, and DAVID 6.8 **Fig. 7b**. ^34,35^.

The immune cell subset was derived from the filtered, integrated Seurat object and included the Macrophage/DC (cluster 10), neutrophil (cluster 19) and lymphocyte (cluster 13) cells. Gene counts were log transformed, scaled and principle component analysis performed as for the integrated object above. UMAP resolution was set to 0.4, which yielded 14 clusters. The clusters were manually assigned based on inspection of DEGs for each cluster, and cells grouped if canonical markers were biologically redundant. We confirmed manual labeling with an automated labeling program in R, SingleR ^36^.

### Analysis of regulons and their activity in the integrated single cell dataset

SCENIC analysis ^37^ was performed using the default setting and mm9-500bp-upstream-7species.mc9nr.feather database was used for data display.

### Pseudotemporal ordering of single cells

We performed pseudotime analysis on the integrated Seurat object containing all cell types as well as the immune cell subset. Cells from each of the seven time points were included and were split into individual gene expression data files organized by previously defined cell type. These individual datasets were analyzed separately through the R package Monocle using default parameters. Outputs were obtained detailing the pseudotime cell distributions for each cell type. Positional information for the monocle plot was used to subset and color cells for downstream analyses ^38^. We performed a separate temporal ordering analysis of S1, S2 and S3 proximal tubule segments across all time points and visualized using t-SNE, produced by Harmony and Palantir R packages ^39^.

### RNA velocity analysis

BAM files were fed through the velocyto pipeline ^40^ to obtain .loom files for each experimental condition. These loom files along with their associated UMAP positions and principal component tables extracted from the merged Seurat file were then fed individually into the RNA Velocity pipeline as described in the Velocyto.R Dentate Gyrus/loom tutorial. The default settings described in the tutorial were used except for tSNE positions that were overwritten with the associated UMAP positions from the merged Seurat object, as well as the principal component table. This generated an RNA velocity Fig. mapped using the merged Seurat object cell positions. Similar analysis was done for the immune subsetted data.

### Cell-cell communication analysis

We applied the Cellphone database ^41^ of known receptor-ligand pairs to assess cell-cell communication in our integrated dataset. Gene expression data from the integrated Seurat file was split by time point and genes renamed to Human gene names then reformatted into the input format described on the CellphoneDB website. Individual time point samples were fed into the web document on the cellphone dB website using 50 iterations, precision of 3, and 0.1 ratio of cells in a cluster expressing a gene. Output files for each time point obtained from the website were merged, then interactions trimmed based on significant sites and only selecting secreted interactions.

To visualize cellular cross talk, we applied this data to a circular plot. The interactions from the merged, trimmed cellphone dB file were sorted by cluster interaction then consolidated into 17 final cell types. Each cell type contained a list of significant interacting pairs (with p < 0.05) and their associated strength values (the larger the value the smaller the p value). These were then visualized using R Circlize package ^42^

### Human sepsis staging

Murine scRNAseq data was pseudobulked though selection of 2000 randomly selected cells for each of seven time points and data normalized using edgeR function calcNormFactors. DEGs were found between one versus rest of time points and significant genes filtered by selecting for FDR <0.05. Human specimens were derived from OCT cores of kidney biopsy or nephrectomy samples (GSE139061). All biopsy specimens (N = 22) had a primary pathology diagnosis of AKI and were acquired in clinical care of patients with a diagnosis of sepsis ^5^. The reference nephrectomies were obtained from unaffected portions of tumor nephrectomies or deceased donors. A bulk 20-µm cross-section was cut from each OCT core and RNA was extracted using the Arcturus Picopure extraction kit (KIT0214, Thermo Fisher Scientific, Waltham, MA). Libraries were prepared with the Takara SMARTer® Stranded Total RNA-Seq Kit v2 - Pico Input. Sequencing was performed on an Illumina HiSeq 4000. The murine genes from each pseudobulk time point were translated to their respective human orthologues using the biomaRt package and ensembl database. Each gene had its expression fold change calculated for each time point in relation to all other time points in the mouse. Separately for each human biopsy specimen, the expression of each gene was calculated as a fold change compared to the mean of all reference samples. A spearman correlation assessed alignment between the fold changes of the mouse and human data. Data were displayed as a heatmap.

### Spatial Transcriptomics

A septic mouse kidney was immediately frozen in Optimal Cutting Temperature media. A 10 µm frozen tissue section was cut and affixed to a Visium Spatial Gene Expression library preparation slide (10X Genomics). The specimen was fixed in methanol and stained with hematoxylin-eosin reagents. Images of hematoxylin-eosin-labeled tissues were collected as mosaics of 10x fields using a Keyence BZ-X810 fluorescence microscope equipped with a Nikon 10X CFI Plan Fluor objective. The tissue was then permeabilized for 12 minutes and RNA was isolated. The cDNA libraries were prepared and then sequenced on an Illumina NovaSeq 6000. Using Seurat 3.1.4, we identified anchors between the integrated single cell object and the spatial transcriptomics datasets and used those to transfer the cluster data from the single cell to the spatial transcriptomics. For each spatial transcriptomics spot, this transfer assigns a score to each single cell cluster. We selected the cluster with the highest score in each spot to represent its single cell associated cluster. Using a Loupe Browser, expression data was visualized overlying the hematoxylin-eosin image.

### Single-molecule RNA in situ hybridization

Formalin-fixed paraffin-embedded cross sections were prepared with a thickness of 5µm. The slides were baked for 60 minutes at 60 °C. Tissues were incubated with Xylene for 5 minutes x2, 100% ETOH for 2 minutes x2, and dried at room temperature. RNA in situ hybridization was performed using RNAscope multiplex Fluorescent Reagent Kit v2 (Advance Cell Diagnosis Inc.) as per the manufacturer instructions. Probe sets were obtained from Advance Cell Diagnosis Inc (murine Agt Cat. No. 426941, Aqp1 Cat. No. 504741-C2). TSA Cyanine 3 Plus and Fluorescein Plus Evaluation kit (PerkinElmer, Inc) was used as secondary probes for the detection of RNA signals. All slides were counterstained with DAPI and coverslips were mounted using fluorescent mounting media (ProLong Gold Antifade Reagent, Life Technologies). The images were collected with a LSM800 confocal microscope (Carl Zeiss).

### Quantification and Statistical Analysis

No blinding was used for animal experiments. All data were analyzed using R software packages, with relevant statistics described in results, methods and Fig. legends.

### Data availability

Data will be deposited to NCBI GEO. The authors declare that all relevant data supporting the findings of this study are available on request.

### Code availability

R scripts for performing the main steps of analysis are available from the Lead contact on request.

## Additional Information

Correspondence and requests for resources and reagents should be directed to and will be fulfilled by the Lead Contact Takashi Hato.

### Supplemental items

Supplemental Fig. 1-7: refer to “Supplemental_Fig 1-7.pdf”

Supplemental Table 1: Cell-type specific differentially expressed genes from 0-48 hours, related to Fig. 1, Supplemental Fig. 1.

Supplemental Table 2: Numbered interactions of receptor ligand interactions between cell type pairs, related to Fig. 6 and Supplemental Fig. 7.

Supplemental Table 3: Scenic regulatory gene network analysis of select cell types highlighting upregulated genes from 16-hour time point, related to Fig. 7.

Supplemental Table 4: Murine pseudo bulked genes and counts from 0-48 hours, related to Fig. 7.

Supplemental Table 5: Human gene count matrices and clinical data from AKI renal biopsies, related to Fig. 7.

## Acknowledgements

We thank the Kidney Precision Medicine Project for making data available for human kidney reference nephrectomy specimens. We thank Daria Barwinska for assistance with specimen validation. This work was supported by NIH K08-DK113223 to TH, NIH R01-DK080063, Veterans Affairs Merit (1I01BX002901) and the Indiana Clinical and Translational Sciences Institute (UL1TR002529) to PCD, K08-DK107864 to MTE, T32HL091816 and T32DK120524 to DJ.

## Author contributions

Conceptualization DJ, PCD and TH. scRNAseq Methodology DJ, BM, PCD and TH. scRNAseq Software, formal analysis, visualization TH, JM, DJ. Investigation DJ, AZ, TH. Validation KC and SW, TA. Resources for single cell data, spatial transcriptomics, and human data TH, PD, TA, MTE, RMF. Data curation, DJ, TH, TWM, JM. Writing-original draft, DJ, PD, TH. Supervision, TH and PD. Funding PD, TH, DJ.

---

**Supplementary Figure 1.**
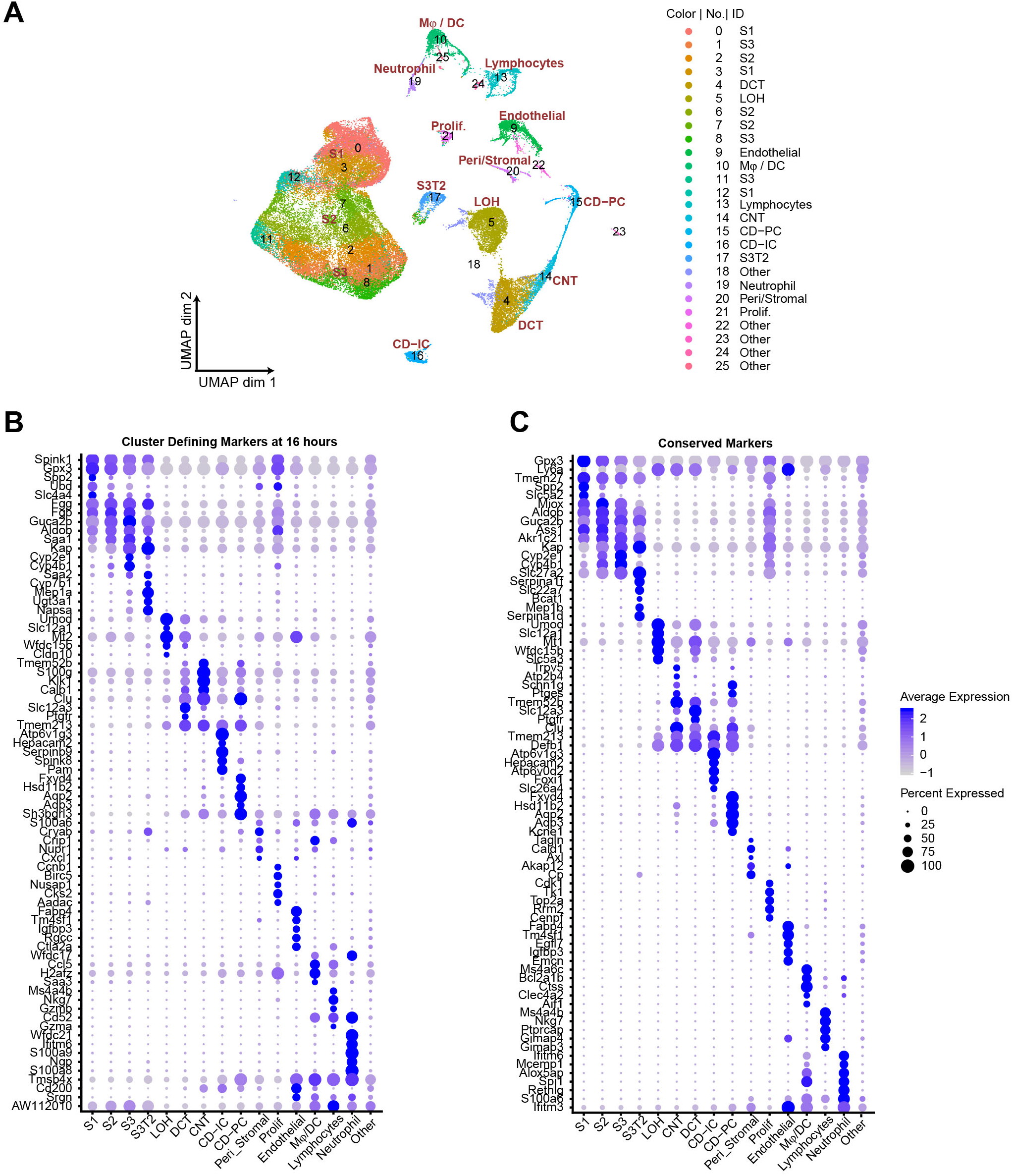
Cluster-defining markers across the sepsis timeline, related to Figure 1. (A) Integrated UMAP of kidney cell clusters showing both assigned identity and original cluster number from control and LPS-treated mice (0, 1, 4, 16, 27, 36 and 48 hours after LPS injection). (B-C) Dot plots of top five cluster-defining (shown at 16 hours) and conserved marker genes (all time points integrated). CD, collecting duct. CD-IC, collecting duct-intercalated cells. CD-PC, collecting duct-principle cells. CNT, connecting tubule. OCT, distal convoluted tubule. Endo, endothelial cells. LOH, Loop of Henle. LPS, endotoxin. Ly, lymphocytes. Mφ-DC, macrophage-dendritic cells. Ne, neutrophil. Peri/St, mixed pericyte and stromal cells. Prolif. Cells, proliferative cells. PT, proximal tubule. S1, first segment of PT. S2, second segment of PT. S3, third segment of PT. S3T2, S3 type 2 cells.

**Supplementary Figure 2.**
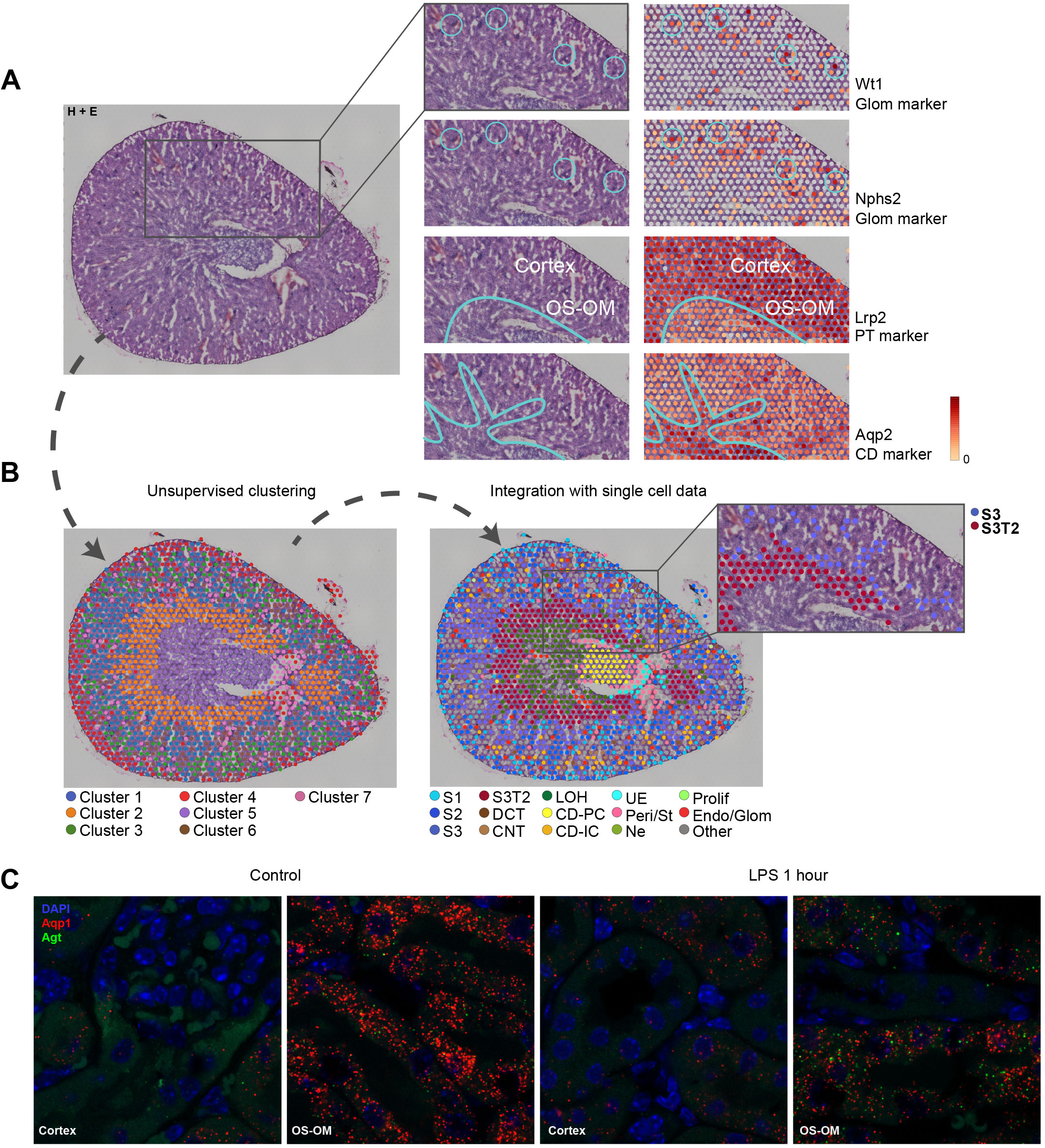
Spatial transcriptomics validation, related to Figure 2. (A-B) Spatial transcriptomics of murine kidney showing unsupervised clustering (B-left) and expanded clustering after integration with single cell data (B-right). Insets of A show gene expression of select glomerular and tubular markers. Inset of B-right highlights S3 and S3T2 clusters. (C) Single molecular FISH (smFISH) coexpression of Aqp1 (red) and Agt (punctate green) in control kidney and after 1 hour of LPS (Cortex and outer stripe of outer medulla shown). Diffuse green objects are RBC and tubular autofluorescence. CD, collecting duct. CD-IC, collecting duct-intercalated cells. CD-PC, collecting duct-principle cells. CNT, connecting tubule. OCT, distal convoluted tubule. Endo/Glom, glomerular endothelial cells. Glom, glomerulus. H+E, hematoxylin and eosin stain, LOH, Loop of Henle. LPS, endotoxin. Ly, lymphocytes. Mφ-DC, macrophage-dendritic cells. Ne, neutrophil. OS-OM, outer stripe of outer medulla. Peri/St, mixed pericyte and stromal cells. Prolif. Cells, proliferative cells. PT, proximal tubule. S1, first segment of PT. S2, second segment of PT. S3, third segment of PT. S3T2, S3 type 2 cells. UE, ureteric epithelium

**Supplementary Figure 3.**
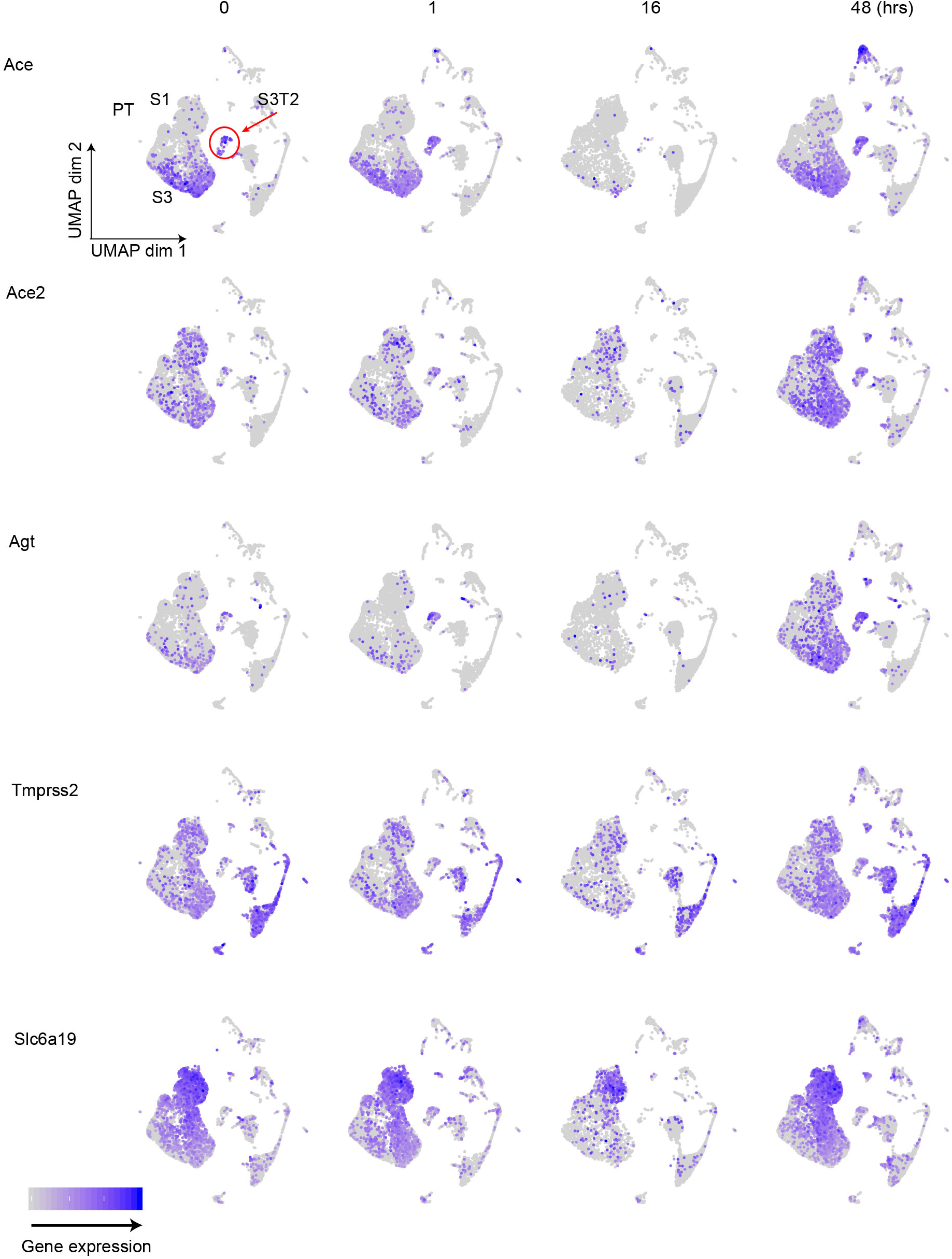
SARS CoV-2 axis, related to Figure 2. Feature plots showing expression of SARS CoV-2 related genes at specified time points. Hr(s), hour(s). PT, proximal tubule. S1, first segment of PT. S3, third segment of PT. S3T2, S3 type 2 cells.

**Supplementary Figure 4.**
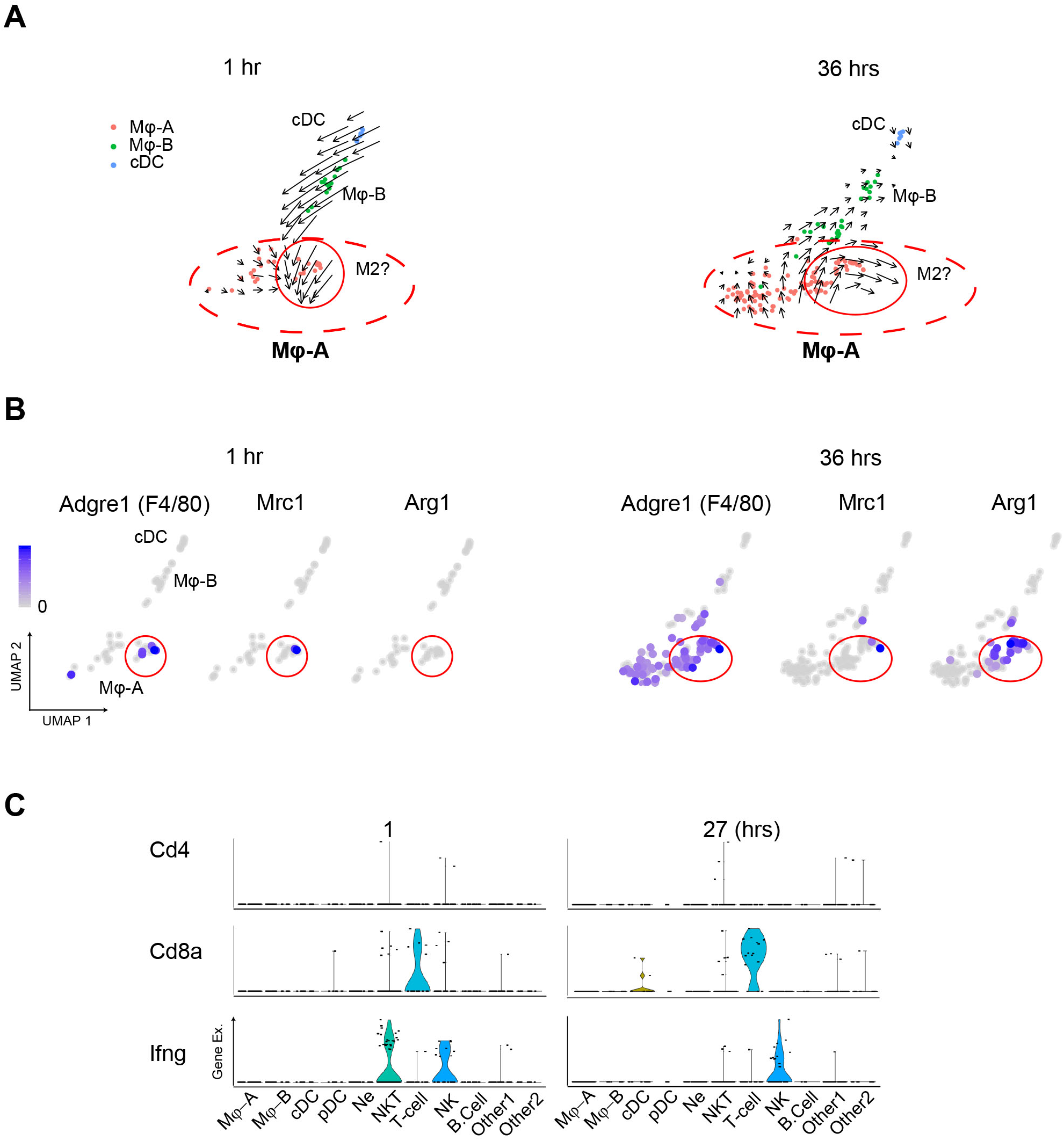
Immune cell subset characteristics, related to Figure 3. (A) RNA velocity analysis reveals two distinct subfields within the Mφ-A cluster. The subfield circled in red showed expression of M2 macrophage-related genes at later time points (B). (C) Violin plots of Cd4, Cd8 and lfn-g expression across immune cell subtypes. cDC, conventional dendritic cell. Hrs, hours. Mφ-A, macrophage-A. Mφ-B, macro-phage-B. M2, alternatively activated macrophages. Ne, neutrophil. NK, natural killer cells. NKT, natural killer T-cells. pDC, plasmacytoid dendritic cell. T-cell, Cd3+ T-lymphocytes.

**Supplementary Figure 5.**
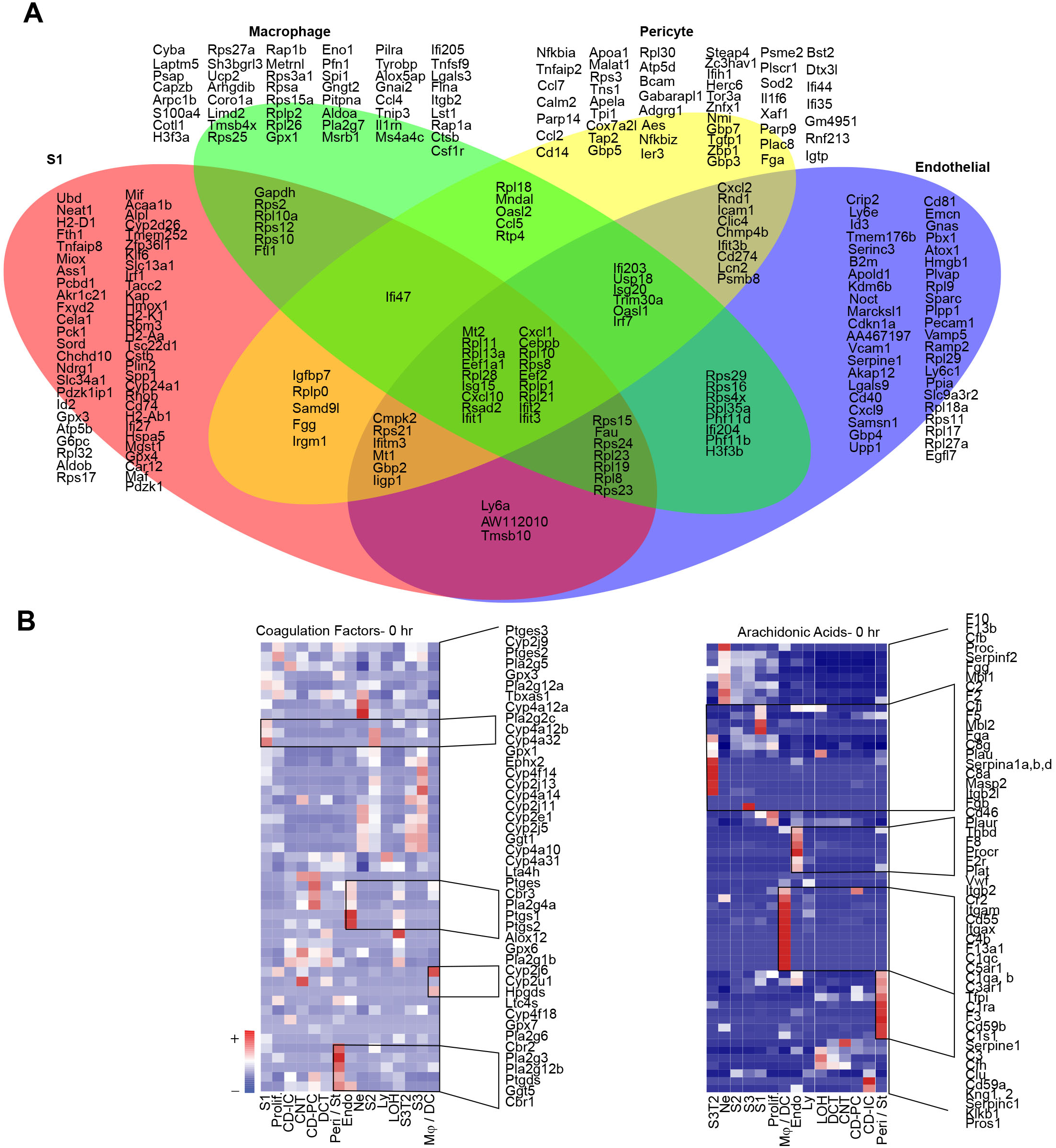
Comparisons of transcriptomic profiles across the sepsis timeline, related to Figure 5. (A) Venn diagram showing differentially expressed genes across time (0-48 hours) for indicated cell types. (B) Heatmaps of genes involved in coagulation and arachidonic acid related pathways in all cell types. CD-IC, collecting duct-intercalated cells. CD-PC, collecting duct-principle cells. CNT, connecting tubule. OCT, distal convoluted tubule. Endo, endothelial cells. Hr(s), hour(s). LOH, Loop of Henle. Ly, lymphocytes. Mφ-DC, macrophage-dendritic cells. Ne, neutrophil. Peri/St, mixed pericyte and stromal cells. Prolif., proliferative cells. PT, proximal tubule. S1, first segment of PT. S2, second segment of PT. S3, third segment of PT. S3T2, S3 type 2 cells.

**Supplementary Figure 6.**
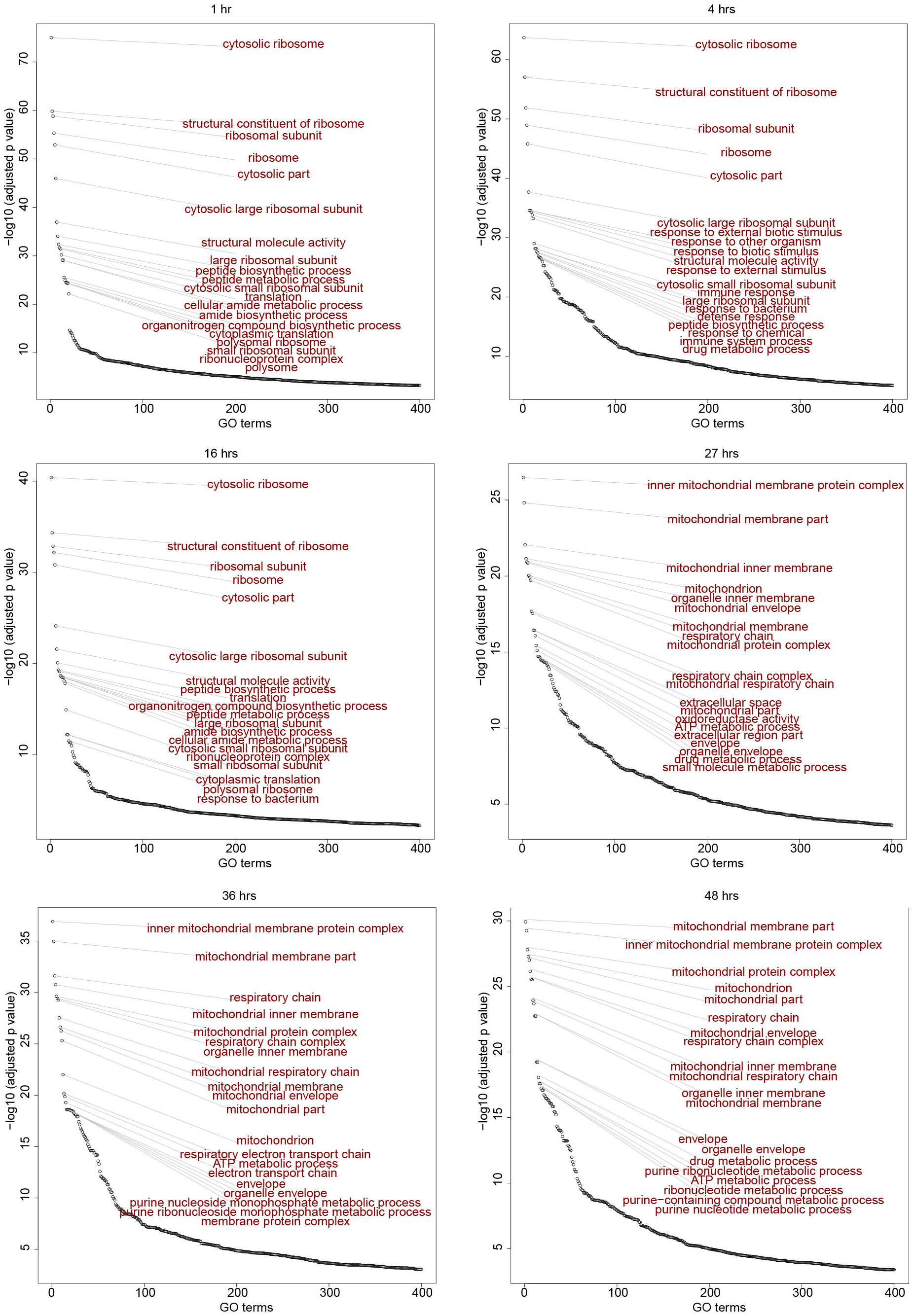
S3 T2 GO terms, related to Figure 5. Time dependent enrichment of gene ontology terms for S3T2 cells. GO terms are sorted in order of statistical significance. Hr(s), hour(s). GO, gene ontology biological processes. S3T2, S3 type 2 cells.

**Supplementary Figure 7.**
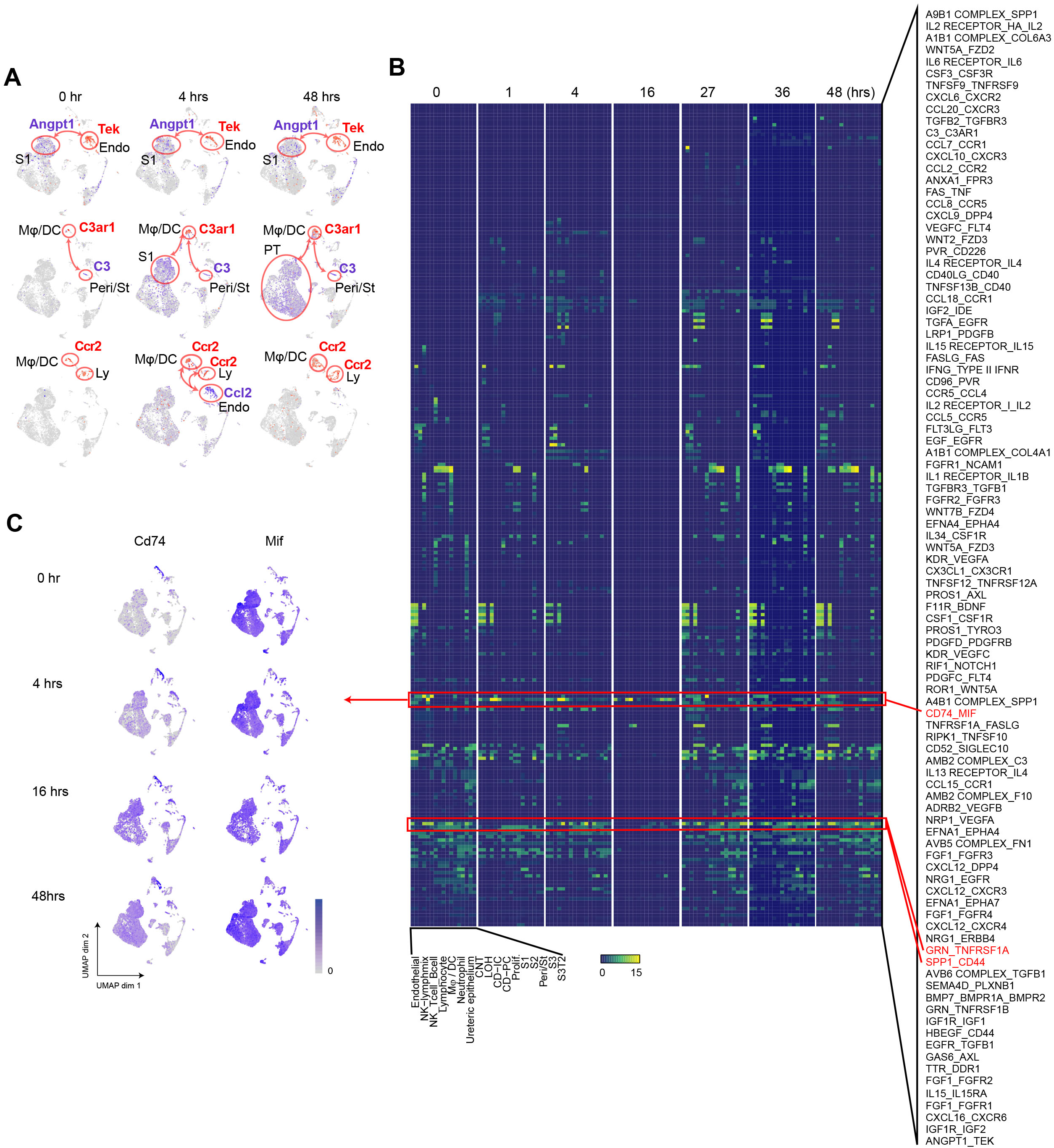
Expanded cell-cell communication examples, related to Figure 6. (A) Feature plots illustrating cell and time-dependent expression changes of select receptor-ligand pairs. (B) Heatmaps of receptor-ligand pairs at select time points. Red boxes highlight select pairs across time. (C) Feature plots illustrating time dependent expression of Cd74-MIF receptor-ligand pair. CD-IC, collecting duct-intercalated cells. CD-PC, collecting duct-principle cells. CNT, connecting tubule. OCT, distal convoluted tubule. Endo, endothelial cells. Hr(s), hour(s). LOH, Loop of Henle. Ly, lymphocytes. Mφ-DC, macro-phage-dendritic cells. Ne, neutrophil. Peri/St, mixed pericyte and stromal cells. Prolif., proliferative cells. PT, proximal tubule. S1, first segment of PT. S2, second segment of PT. S3, third segment of PT. S3T2, S3 type 2 cells.

## Notes

### Competing Interest Statement

The authors have declared no competing interest.

